# Catalytic Promiscuity of Rice 2-Oxoglutarate/Fe(II)-Dependent Dioxygenases Supports Xenobiotic Metabolism

**DOI:** 10.1101/2020.12.17.423306

**Authors:** Natsuki Takamura, Akihiko Yamazaki, Nozomi Sakuma, Sakiko Hirose, Yukie Takani, Motonari Sakai, Masahiro Oshima, Makoto Kuroki, Yuzuru Tozawa

**Affiliations:** Graduate School of Science and Engineering, Saitama University, Saitama 338-8570, Japan; Tsukuba Research & Technology Center, SDS Biotech K.K., Tsukuba 300-2646, Japan; Institute of Agrobiological Sciences, National Agriculture and Food Research Organization, Tsukuba 305-8634, Japan; Institute of Crop Science, National Agriculture and Food Research Organization, Tsukuba 305-8518, Japan

## Abstract

The rice 2-oxoglutarate/Fe(II)–dependent dioxygenase HIS1 mediates the catalytic inactivation of five distinct β-triketone herbicides (bTHs). During a search for potential inhibitors of HIS1, we found that it mediates the hydroxylation of trinexapac-ethyl (TE) in the presence of Fe^2+^ and 2-oxoglutarate. TE is a plant growth regulator that blocks gibberellin biosynthesis, and we observed that its addition to culture medium induced growth retardation of rice seedlings in a concentration-dependent manner. Similar treatment with hydroxylated TE revealed that hydroxylation greatly attenuated the inhibitory effect of TE on plant growth. Forced expression of *HIS1* in a rice *his1* mutant also reduced its sensitivity to TE compared with that of the nontransformant. These results indicated that HIS1 metabolizes TE and thereby markedly reduces its ability to slow plant growth. Furthermore, testing of five HIS1-related proteins (HSLs) of rice revealed that OsHSL2 and OsHSL4 also metabolize TE in vitro. HSLs from wheat and barley also showed such activity. In contrast, OsHSL1, which shares the highest amino acid sequence identity with HIS1 and metabolizes the bTH tefuryltrione, did not manifest TE-metabolizing activity. Site-directed mutagenesis of OsHSL1 informed by structural models showed that substitution of three amino acids with the corresponding residues of HIS1 conferred TE-metabolizing activity similar to that of HIS1. Our results thus reveal a catalytic promiscuity of HIS1 and its related enzymes that supports xenobiotic metabolism in plants.

**One-sentence summary:** The rice 2-oxoglutarate/Fe(II)-dependent dioxygenase HIS1 and related enzymes show broad substrate specificity and mediate metabolism of the growth regulator trinexapac-ethyl as well as of herbicides.

## INTRODUCTION

Modern agriculture essentially depends on the many types of chemicals that are now applied for the control of crop diseases, insect pests, weeds, and plant growth and development. Weed control by herbicides relies on intrinsic or artificially introduced enzymes in the crop that are able to mediate metabolic inactivation of the applied chemicals (Kraehmer et al., 2014a, 2014b).

We previously described the isolation and characterization of a rice herbicide resistance gene, *HIS1*, that encodes an Fe(II)- and 2-oxoglutarate dependent dioxygenase (2OGD) (Maeda et al., 2019). Biochemical analysis and plant transformation tests revealed that HIS1 catalyzes the hydroxylation of five β-triketone herbicides (bTHs): benzobicyclon (BBC), tefuryltrione (TFT), sulcotrione, mesotrione, and tembotrione. These bTHs target 4-hydroxyphenylpyruvate dioxygenase (HPPD), which is structurally unrelated to 2OGD enzymes but mechanistically grouped with α-keto acid–dependent oxygenases and members of the 2OGD superfamily (He and Moran 2009). We also uncovered wide conservation of genes that encode HIS1-like (HSL) proteins in other major crops including wheat, corn, barley, and sorghum. A search of the genome database for rice *(Oryza sativa* L. cv. Nipponbare) revealed five predicted HSLs in addition to HIS1. We further found that rice OsHSL1 metabolizes TFT and that forced expression of *OsHSL1* conferred TFT resistance in rice and *Arabidopsis* (Maeda et al., 2019), whereas other HSLs did not manifest metabolic activity for bTHs. The comprehensive conservation of *HSL* genes among major crops was suggestive of common and important functions, but the natural substrates of the encoded enzymes have remained unknown.

All the predicted HSL proteins possess conserved signature motifs of 2OGD enzymes in their amino acid sequences (Hegg and Que, 1997; Wilmouth et al., 2002; Kawai et al., 2014). 2OGD enzymes rely on 2-oxoglutarate (2OG) as a cosubstrate and convert it to succinate. Several compounds that mimic 2OG or succinate and impede the enzyme reaction have been developed as gibberellin biosynthesis inhibitors (Brown et al., 1997; Rademacher, 2000; Ervin and Koski, 2001; McCarty et al., 2004; Rademacher, 2016). Some of these compounds are applied as growth retardants for turfgrass management (Rademacher, 2000; Baldwin et al., 2006; Xu et al., 2016; Rademacher, 2016). On the other hand, the target enzymes of these growth retardants have not been specified because of their potentially broad spectrum of action. We initially conjectured that these compounds might also affect HSL enzymes, and we began to test general 2OGD inhibitors—such as daminozide, prohexadione, and trinexapac-ethyl (TE)—for potential inhibitory effects on HIS1 activity in vitro. During this testing, we found that the rice enzyme HIS1 metabolizes TE and we confirmed that the metabolite produced by HIS1 is a hydroxylated form of TE (TE-OH). We further observed that forced expression of *HIS1* reduces the TE sensitivity of rice. OsHSL1, the closest paralog of HIS1, was found not to possess TE-hydroxylating activity, whereas OsHSL2 and OsHSL4, both of which are less similar to HIS1 than is OsHSL1, did manifest HIS1-like TE-metabolizing activity. Mutation analysis based on a comparison of the amino acid sequences of HIS1 and OsHSL proteins revealed that the substitution of three amino acids of OsHSL1 with the corresponding residues of HIS1 conferred TE-metabolizing activity. In addition, HSL proteins of wheat and barley were shown to possess TE-metabolizing activity in vitro. These findings have thus revealed two specific characteristics of HIS1 and HSL enzymes: (1) their catalytic activities do not necessarily reflect amino acid sequence similarity, and (2) their substrates include a broad-spectrum 2OGD inhibitor.

## RESULTS

### Prohexadione Partially Inhibits the bTH-Metabolizing Activity of HIS1 in Vitro

We previously showed that *HIS1*-like (*HSL*) genes are highly conserved among poaceous species, suggestive of common and important roles of *HSL* gene products in major crops (Maeda et al., 2019). However, the natural substrate (or substrates) of HIS1 and its related proteins have remained to be identified. To determine conditions that allow accumulation of the natural substrate of HIS1 in plants, we explored the potential inhibitory effects of three general 2OGD inhibitors—daminozide, prohexadione, and TE—on HIS1 activity in vitro (Fig. 1A). For these assays, we used benzobicyclon hydrolysate (BBC-OH) (Fig. 1A) as a substrate of HIS1, as described previously (Maeda et al., 2019).

**Figure 1.**
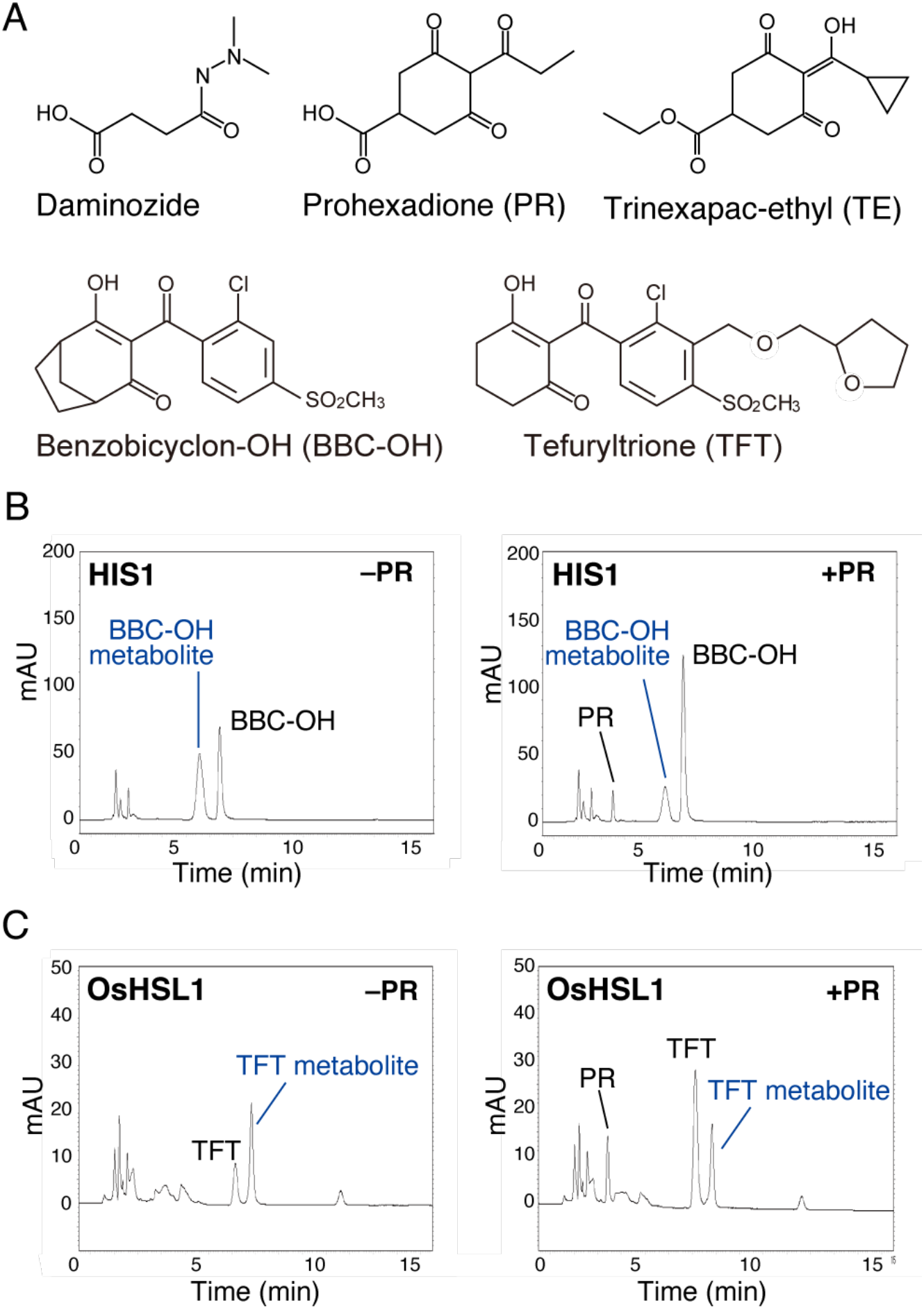
Test for potential inhibitors of HIS1 and OsHSL1 enzyme activity in vitro. A, Structures of chemical compounds used in the study. B, HPLC analysis of reaction mixtures after assay of HIS1 activity with BBC-OH as substrate in the absence or presence of prohexadione (50 μM). Peak positions for BBC-OH, the BBC-OH metabolite (hydroxylated form), and prohexadione are indicated. mAU, milliabsorbance unit. C, HPLC analysis of reaction mixtures after assay of OsHSL1 activity with TFT as a substrate in the absence or presence of prohexadione (50 μM). Peak positions for TFT, the TFT metabolite, and prohexadione are indicated.

We found that prohexadione partially inhibited the BBC-OH hydroxylation activity of recombinant HIS1 in vitro (Fig. 1B), whereas daminozide showed no effect on the enzyme activity (data not shown). Given that the amino acid sequence of OsHSL1 is 87% identical to that of HIS1, we next examined the effect of prohexadione in an OsHSL1 enzyme assay with TFT (Fig. 1A) as the substrate. Prohexadione also partially inhibited the TFT hydroxylation activity of OsHSL1 (Fig. 1C). The median inhibitory concentration (IC_50_) for the effect of prohexadione on HIS1 activity was relatively high at 50 μM (Supplemental Fig. S1), and its efficacy was likely not sufficient for suppression of HIS1 and OsHSL1 catalysis in plants.

### HIS1 Metabolizes TE in Vitro

We next tested the effect of TE on HIS1 catalysis with BBC-OH as substrate. Unexpectedly, we found that TE is recognized as a substrate by HIS1 (Fig. 2A). In the reaction catalyzed by HIS1 in the presence of both Fe^2+^ and 2OG, TE was converted to two metabolites represented by a dominant peak (metabolite 1) and a smaller peak (metabolite 2) in the HPLC profile. This conversion did not occur in the absence of either Fe^2+^ or 2OG (Fig. 2B), indicating that it was attributable to the function of HIS1 as a 2OGD.

**Figure 2.**
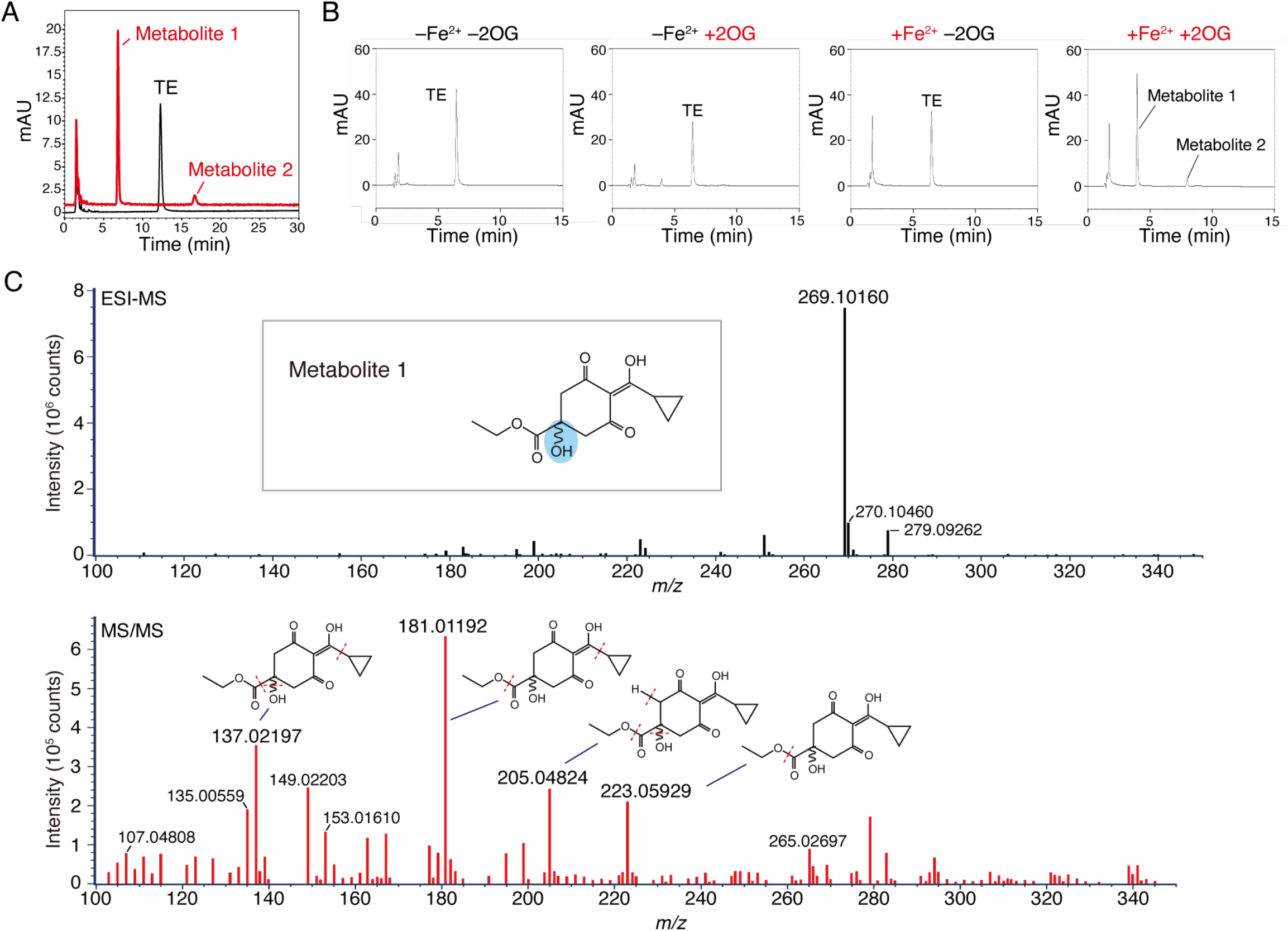
Hydroxylation of TE by HIS1 in vitro. A, LC profile of TE and its HIS1-generated metabolites prior to ESI-MS analysis. The red trace indicates the profile of the reaction mixture after incubation of recombinant HIS1 with TE in the presence Fe^2+^ and 2OG. As a negative control, recombinant green fluorescent protein (GFP) was tested and profiled under equivalent conditions (black trace). B, HPLC profile of the reaction mixture after incubation of HIS1 with TE in the absence or presence of Fe^2+^ or 2OG as indicated. The chromatography conditions differ from those in (A) and are described in Materials and Methods. C, MS analysis and predicted structure of TE metabolite 1 generated by HIS1 catalysis. Data from LC-ESI-MS analysis (upper panel) and MS/MS analysis (lower panel) in the positive mode are shown for purified metabolite 1 as in (A). The predicted structure of the compound based on these results and NMR data is also shown. The blue shading indicates the hydroxylated position. Predicted cleavage positions for metabolite 1 in the MS/MS analysis are indicated by red dotted lines.

We isolated these two TE metabolites and analyzed their structures. Analysis by liquid chromatography and electrospray-ionization mass spectrometry (LC-ESI-MS) revealed that metabolite 1 (mass charge ratio *[m/z]* = 269.10160) (Fig. 2C) has a mass that is greater than that of the parent molecule *(m/z* = 253.11144) (Supplemental Fig. S2A) by 16. In contrast, the mass of metabolite 2 *(m/z* = 251.09056) was found to be smaller than that of the parent molecule (Supplemental Fig. S2B). NMR analysis of the isolated metabolite 1 (hereafter referred to as TE-OH) revealed that it possesses a hydroxyl group at the *p*-position (Supplemental Table S1). We therefore identified TE metabolite 1 as ethyl 4-(cyclopropanecarbonyl)-1-hydroxy-3,5-dioxocyclohexane-1-carboxylate (Fig. 2C). We predicted metabolite 2 to be a cyclohexene derivative (Supplemental Fig. S2B) on the basis of the ESI-MS and tandem MS (MS/MS) data.

### In Vivo Effects of TE and TE-OH in Rice

We previously identified two rice *his1* mutants that lack a functional HIS1 enzyme as a result of Tos17 retrotransposon insertion in an exon at the *HIS1* locus (Maeda et al., 2019). We therefore performed a growth test for *his1* mutant seedlings and Nipponbare with or without TE treatment. We evaluated elongation of the second leaf sheath of seedlings as a measure of gibberellin action. Treatment of Nipponbare with TE resulted in a smaller second leaf sheath length compared with that of untreated seedlings, with this effect of TE being concentration dependent (Fig. 3A). We obtained similar results with the two *his1* mutants (data not shown), suggesting that the effect of TE on seedling growth is not influenced by the lack of HIS1.

**Figure 3.**
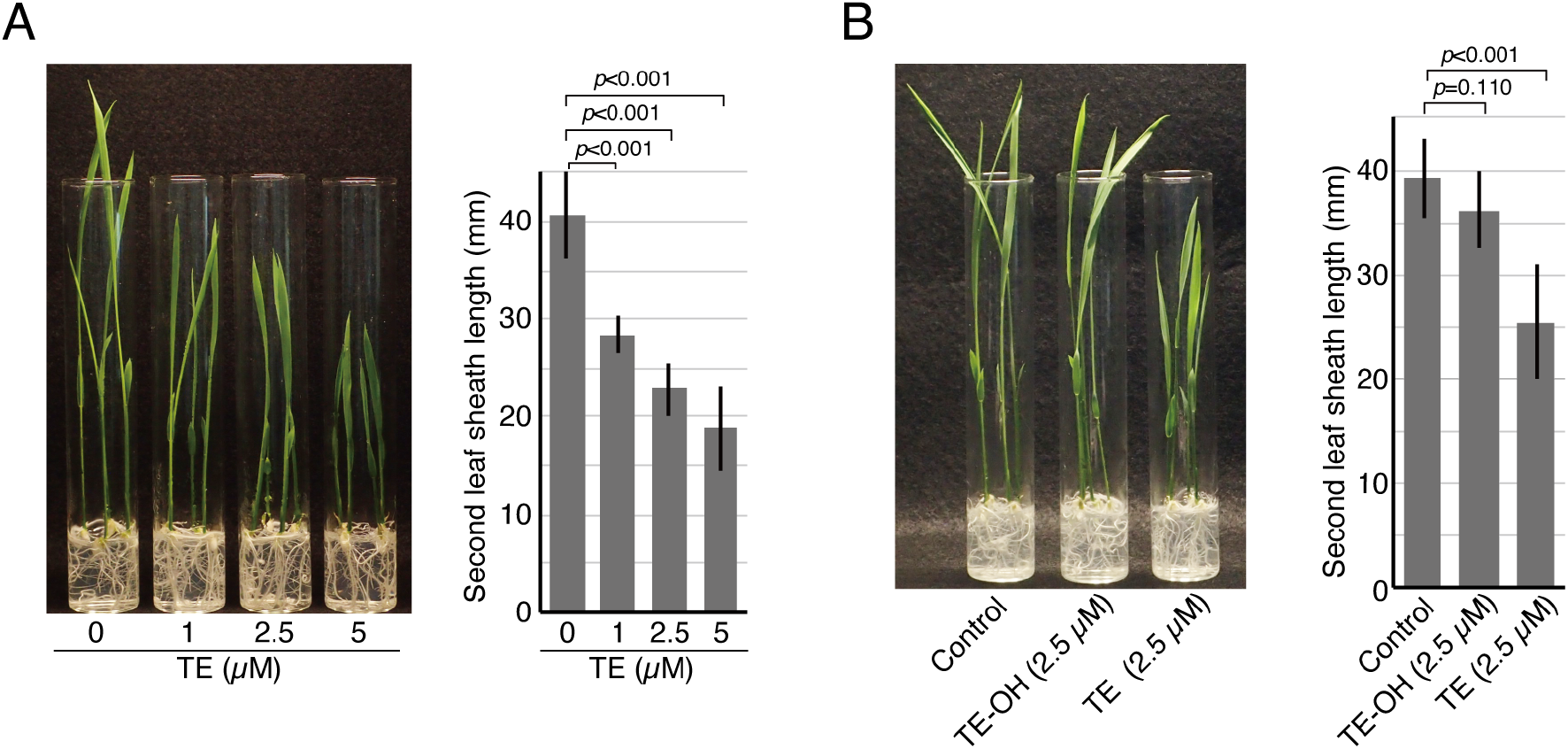
Effects of TE and TE-OH on the growth of rice seedlings. Seeds of rice cultivar Nipponbare were germinated and cultured on solid medium for 7 days in the absence or presence of the indicated concentrations of TE (A) or of TE or TE-OH (B). Representative seedlings as well as quantitative data for the length of the second leaf sheath are shown. The quantitative results are mean ± SD values for 11 to 13 independent biological replicates (see Supplemental Table S2). The levels of statistical significance are shown as *p*-values based on evaluation by Student’s t test.

Given that HIS1 was shown to catalyze the hydroxylation of TE in vitro (Fig. 2), we next examined whether such hydroxylation might abolish the toxicity of TE by testing the effect of TE-OH on the growth of rice seedlings. TE was converted to TE-OH by the HIS1 enzyme reaction, and the effects of TE-OH and TE on seedling growth were then compared. In contrast to the inhibitory effect of TE on internode elongation in Nipponbare seedlings, TE-OH had a much weaker inhibitory effect (Fig. 3B, Supplemental Table S2), indicating that hydroxylation of TE by HIS1 greatly diminishes its inhibitory effect on plant growth.

We next tested the TE sensitivity of rice strains that overexpress *HIS1* under the control of the 35S promoter of cauliflower mosaic virus (Maeda et al., 2019). Forced expression of *HIS1* rendered Yamadawara, which is a *his1/his1* mutant homozygote, partially tolerant to TE. The second leaf sheath of the *HIS1*-overexpressing seedlings was thus longer than that of the parental strain after TE treatment (Fig. 4, Supplemental Table S3). The transformant line #6 appeared to be more resistant to TE than was transformant line #5. Given that we previously showed that the *HIS1* expression level in line #6 is higher than that in line #5 (Maeda, 2019), the TE insensitivity phenotype of the transformant lines appeared to reflect the level of *HIS1* expression.

**Figure 4.**
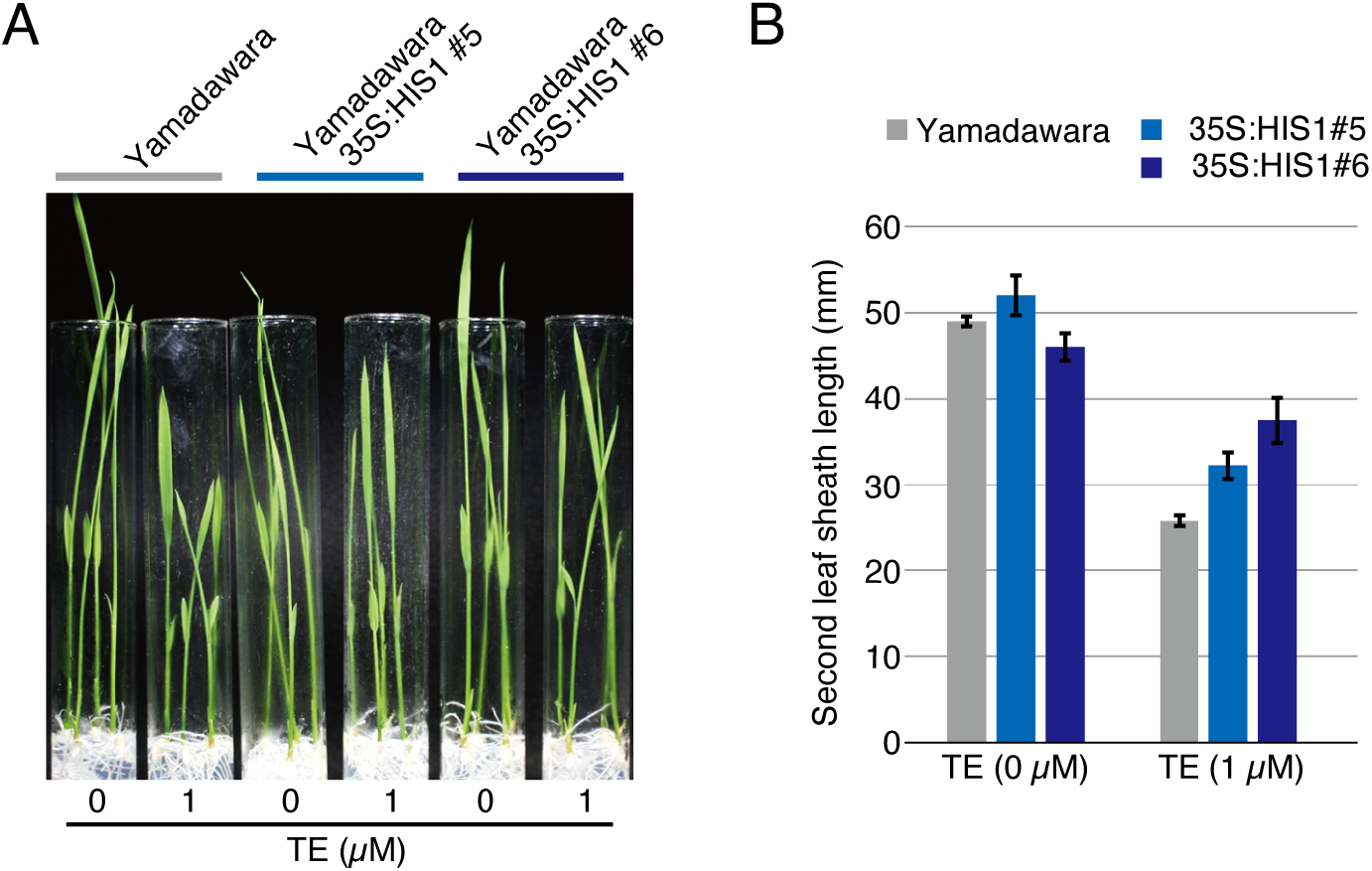
Effect of *HIS1* overexpression on TE sensitivity in a *his1* rice mutant line. A, Seeds of Yamadawara *(his1* mutant) or its transformant lines #5 and #6 that express *HIS1* under the control of the 35S promoter of cauliflower mosaic virus were germinated and cultured on solid medium for 7 days in the absence or presence of 1 μM TE. B, Length of the second leaf sheath for seedlings as in (A). Data are means ± SD for three or four independent biological replicates (see Supplemental Table S3). The levels of statistical significance are shown as *p*-values based on evaluation by Student’s t test.

### OsHSL2, OsHSL4, and Other HSL Proteins Possess TE-Metabolizing Activity in Vitro

We previously showed that HIS1 and OsHSL1 share 87% amino acid sequence identity and that HIS1 catalyzes the hydroxylation of multiple bTHs, including BBC-OH and TFT, whereas OsHSL1 catalyzes the hydroxylation only of TFT (Maeda et al., 2019). We next performed the TE hydroxylation assay with OsHSL1 and other HSL proteins including OsHSL2, OsHSL4, OsHSL5, and OsHSL6 *(O. sativa* cv. Nipponbare); TaHSL1 and TaHSL2 *(Triticum aestivum);* HvHSL1A, HvHSL6A, and HvHSL6D *(Hordeum vulgare*); SbHSL1 *(Sorghum bicolor);* and ZmHSL1A and ZmHSL1B (*Zea mays)* (Fig. 5, A and B). We found that OsHSL1 did not manifest TE conversion activity (Fig. 5C). Further examination of the effect of TE on OsHSL1 function revealed that it slightly inhibited OsHSL1 activity with a relatively high IC_50_ of ~125 μM for TFT conversion (Supplemental Fig. S3). On the other hand, the two rice enzymes OsHSL2 and OsHSL4, the two wheat enzymes TaHSL1 and TaHSL2, and the barley enzyme HvHSL6A all showed TE hydroxylation activity in vitro (Fig. 5C, Table 1). The TE metabolites generated in the reactions with OsHSL4 and HvHSL6A showed different HPLC profiles compared with those generated by the other enzymes (Fig. 5C). The other HSL proteins tested did not show TE-metabolizing activity (Supplemental Fig. S4). The observed activity for TE metabolism of these various HSL proteins did not appear to reflect the phylogenetic relations among the proteins (Fig. 5A), suggesting that overall similarity in amino acid sequence does not necessarily correspond to that in catalytic function in TE metabolism.

**Figure 5.**
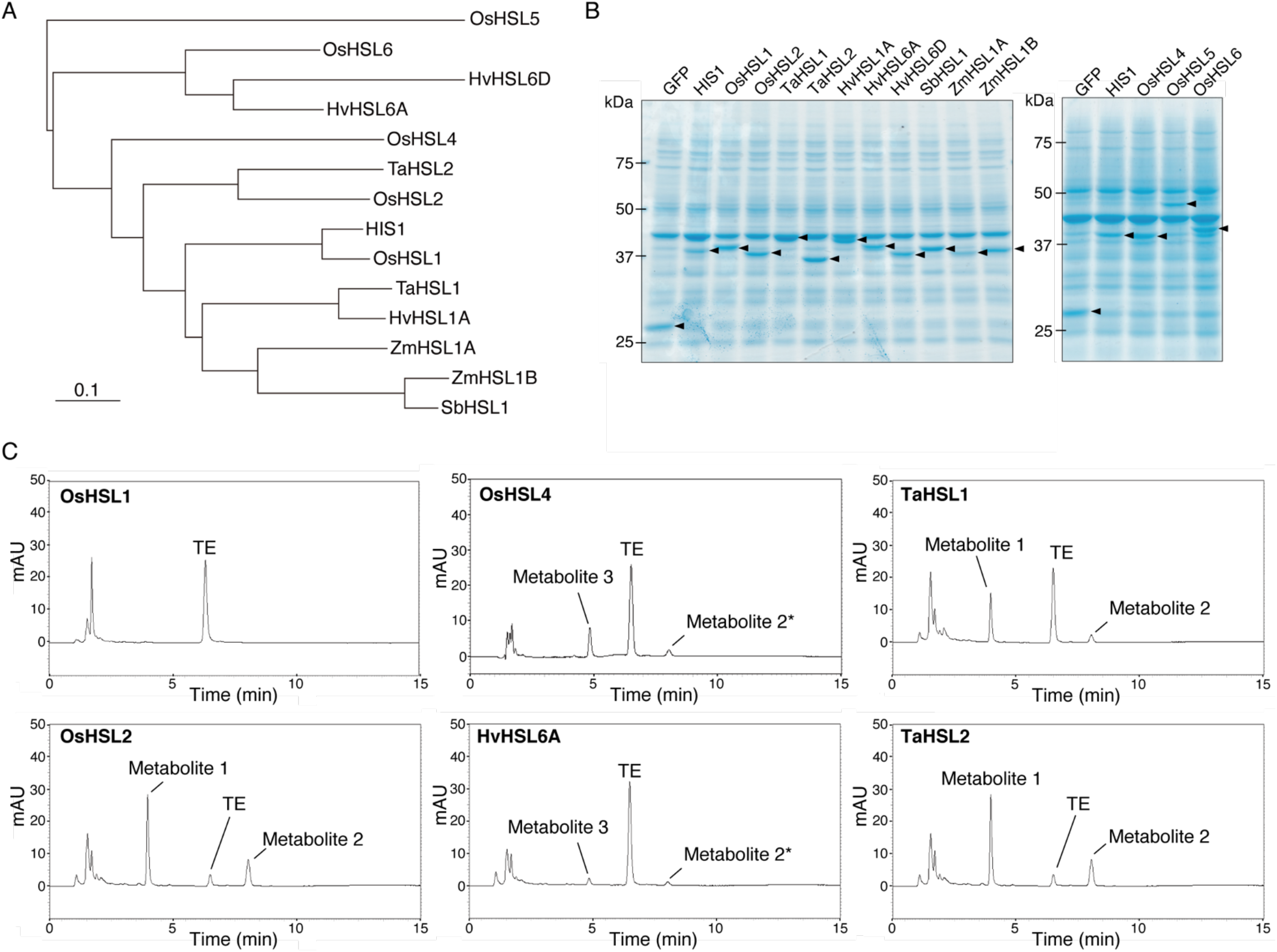
TE-metabolizing activity of various HSL proteins in vitro. A, Phylogenetic tree for HIS1 and HSL proteins analyzed in this study. Prefixes Os, Ta, Hv, Sb, and Zm indicate *Oryza sativa, Triticum aestivum, Hordeum vulgare, Sorghum bicolor,* and *Zea mays,* respectively. The scale bar indicates 0.1 amino acid changes per alignment position. B, Coomassie brilliant blue–stained gel after SDS-PAGE of HIS1, HSL proteins, and GFP (control) produced in a cell-free translation system. Arrowheads indicate the specific recombinant proteins. Sizes (kDa) of molecular mass markers are indicated on the left. C, HPLC profiles of the reaction mixtures after incubation of recombinant OsHSL1, OsHSL2, OsHSL4, HvHSL6A, TaHSL1, or TaHSL2 with TE in the presence of Fe^2+^ and 2OG. Each experiment was performed at least twice to confirm reproducibility.

**Table 1:**
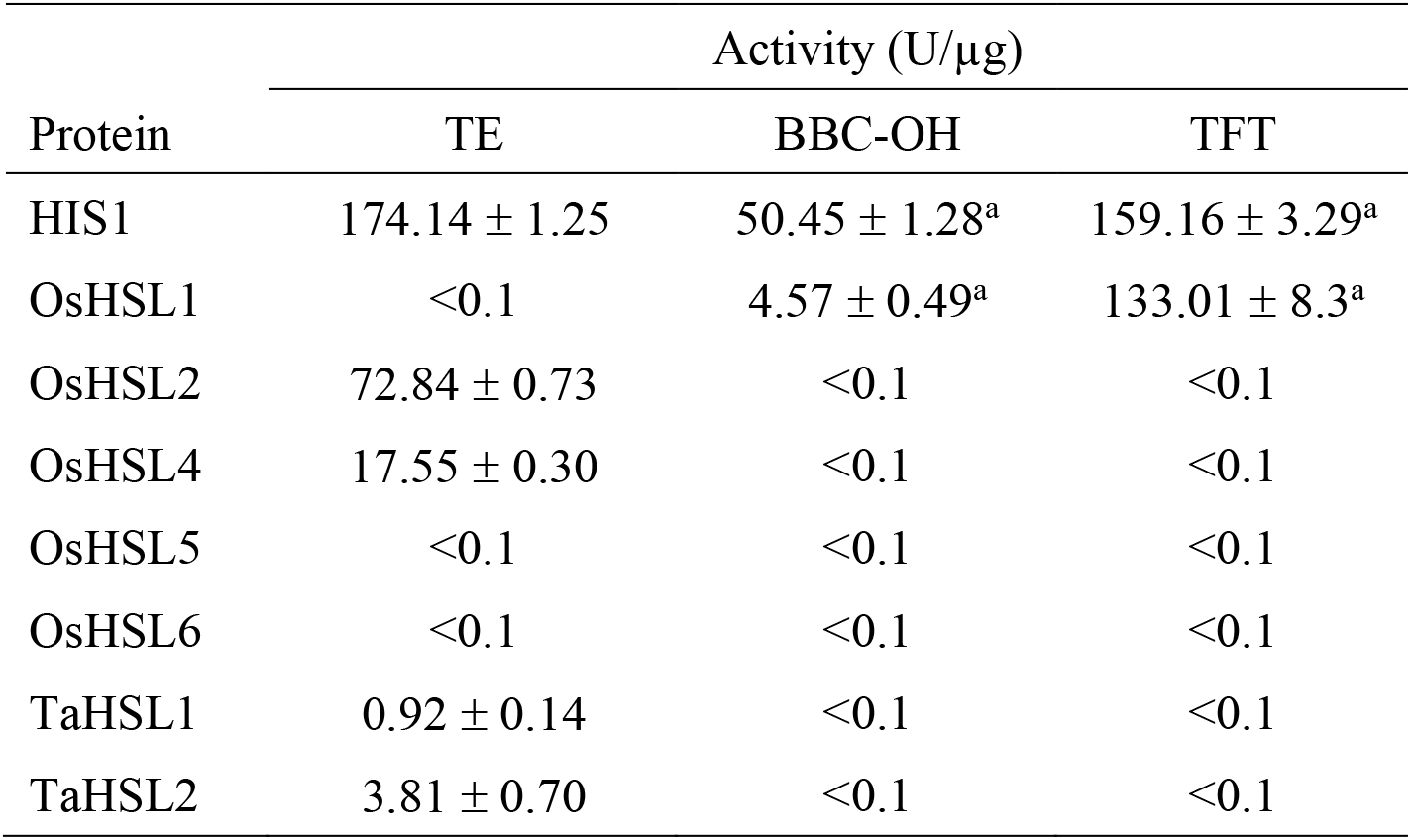
Metabolic activities of HIS1 and various HSL proteins for TE, BBC-OH, and TFT.

### Amino Acid Substitutions in OsHSL1 Confer TE-Metabolizing Activity

We next carried out mutation analysis for OsHSL1 based on comparison of its amino acid sequence with those of HIS1 and OsHSL2. Mutant OsHSL1 enzymes were synthesized in a cell-free system and assayed for their ability to metabolize TE in vitro. We focused on the putative substrate-recognition pocket of OsHSL1—in particular, on the three residues Phe^140^, Leu^204^, and Phe^298^—for mutation analysis (Fig. 6A). Single amino acid substitutions based on the HIS1 sequence—OsHSL1_F140H, OsHSL1_L204F, and OsHSL1_F298L—had essentially no effect on the catalytic activity of OsHSL1 with TE as substrate (Supplementary Fig. S5). In contrast, the double-mutants OsHSL1_F140H/L204F and OsHSL1_F140H/F298L showed substantial TE-metabolizing activity (Fig. 6B), although OsHSL1_L204F/F298L possessed minimal activity (Supplemental Fig. S5). Furthermore, the combination of all three mutations (OsHSL1_F140H/L204F/F298L) conferred a level of TE-metabolizing activity similar to that of HIS1 (Fig. 6B, Supplemental Fig. S5).

**Figure 6.**
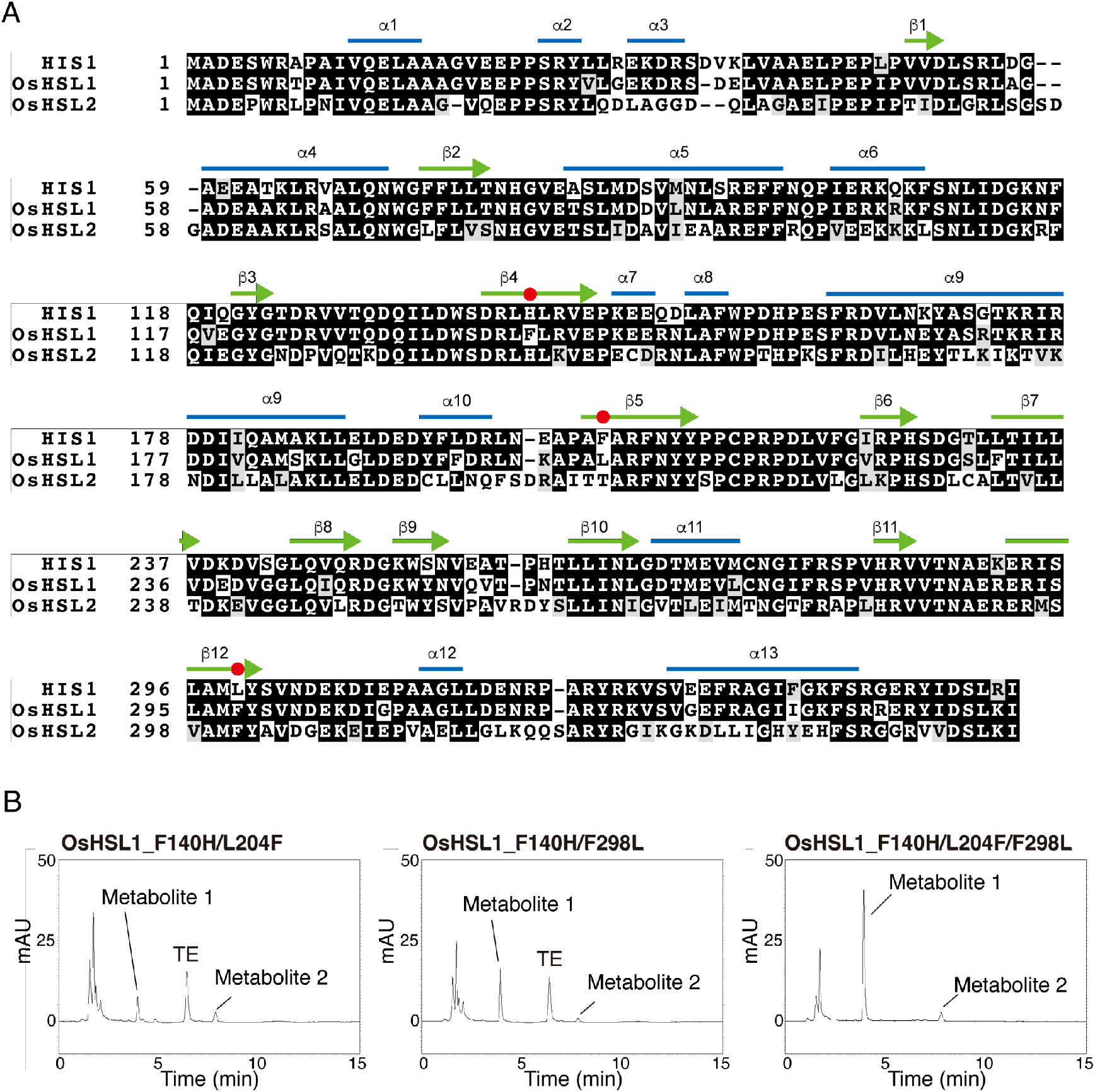
Mutation of specific residues of OsHSL1 confers TE-metabolizing activity. A, Alignment of the predicted amino acid sequences of HIS1 (GenBank accession no. BCK50777), OsHSL1 (BCK50778), and OsHSL2 (LC505696) from *O. sativa*. The alignment was performed with the CLUSTALW algorithm. Blue lines and green arrows indicate predicted a-helix and β-sheet secondary structures, respectively. Red circles indicate the positions of amino acid substitutions in OsHSL1. B, HPLC profiles of the reaction mixtures after incubation of the OsHSL1 mutant proteins OsHSL1_F140H/L204F, OsHSL1_F140H/F298L, or OsHSL1_F140H/L204F/F298L with TE in the presence of Fe^2+^ and 2OG. Each assay was performed at least twice to confirm reproducibility.

## DISCUSSION

The present study originated from tests of the effects of three general 2OGD inhibitors—daminozide, prohexadione, and TE—on HIS1 enzyme activity. All three of these compounds act as late-stage gibberellin biosynthetic inhibitors (Rademacher, 2000; Rademacher, 2016). We found that prohexadione partially inhibited the activities of both HIS1 and OsHSL1 in vitro. TE also had a relatively weak inhibitory effect on OsHSL1 activity. These compounds were designed as 2OG- or succinate-mimicking compounds and are therefore could affect multiple 2OGD enzymes (Rademacher, 2000; Rademacher, 2016). Our results now indicate that these general 2OGD inhibitors indeed affect some members of the HSL family of enzymes in plants. However, their inhibitory effects on HIS1 or OsHSL1 activity appear to be insufficient to result in a substantial disturbance of cellular metabolism mediated by these enzymes.

On the other hand, we unexpectedly found that HIS1 metabolizes TE in vitro. As far as we are aware, this is the first identification of a TE-metabolizing enzyme in plants. In agricultural practice, TE is applied worldwide as an antilodging agent for cereals (Rademacher and Bucci, 2002; Rademacher, 2016) and a chemical ripener for sugarcane (van Heerden, 2014). It is also used for turf management (Fagerness et al., 2002; McCann and Huang, 2007). Our finding that HIS1 metabolizes TE may contribute to further studies on the plant species–specific metabolism of TE.

We confirmed that the metabolism of TE by HIS1 is dependent on Fe^2+^ and 2OG, showing that the reaction occurs by the typical mode of 2OGD action (Wilmouth et al., 2002). Furthermore, NMR and MS analyses revealed that HIS1 catalyzes the hydroxylation of TE, resulting in the formation of ethyl 4-(cyclopropanecarbonyl)-1-hydroxy-3,5-dioxocyclohexane-1-carboxylate (TE-OH) as the primary product.

We prepared TE-OH by HIS1-mediated conversion of TE in vitro and administered the TE-OH aqueous solution in seedling growth tests. The exogenously applied TE-OH had a much weaker inhibitory effect on rice seedling growth compared with TE. However, the ability of seedlings to take up TE-OH is presently unknown. On the other hand, our results with *HIS1*-overexpressing rice lines indicated that the conversion of TE to TE-OH by HIS1 attenuates the inhibitory effect of TE on seedling growth, suggesting that this conversion partially protects plants from the inhibitory effect of TE on the gibberellin biosynthetic pathway. However, it remains to be determined whether TE-OH indeed lacks an inhibitory effect on gibberellin biosynthesis.

Cytochrome P450 (CYP) enzymes play pivotal roles in drug metabolism in humans (Anzenbacher and Anzenbacherová, 2001). In particular, CYP3A4 is the most abundant CYP in the liver and metabolizes drugs administered orally with a broad substrate specificity (Brown et al., 2008; Lokwani et al., 2020). CYPs have also been found to play a role in xenobiotic metabolism in plants (Werck-Reichhart et al., 2000; Siminszky, 2006; Iwakami et al., 2014; Saika et al., 2014). However, CYPs also play important roles in secondary metabolism in plants and appear to have broad substrate specificity (Mizutani and Ohta, 2010; Nelson and Werck-Reichhart, 2011). The potential catalytic promiscuity of plant CYPs is likely controlled by the formation of metabolons with other related enzymes in specific metabolic pathways such as flavone biosynthesis (Nakayama et al., 2019). In addition to CYP enzymes, 2OGD proteins also form a large enzyme superfamily in plants (Farrow and Facchini, 2014; Kawai et al., 2014). In the rice genome sequence database, 334 CYP and 114 2OGD genes have been annotated (Kawai et al., 2014). As an example of the broad substrate specificity of 2OGD enzymes, we previously showed that rice HIS1 hydroxylates at least five different bTHs and abolishes their herbicidal function (Maeda et al., 2019). The *HIS1* gene is therefore a resource as a multiple-herbicide resistance gene for the breeding of rice and other crops. We have now revealed that HIS1 also hydroxylates the plant growth regulator TE and thereby attenuates its biological activity. This single enzyme is thus now known to metabolize a total of six different compounds.

With regard to TE metabolism in rice, we further showed that the enzymes OsHSL2 and OsHSL4 also possess TE-metabolizing activity. However, these two HSL proteins have no bTH-inactivating activity. We also found that two wheat HSL proteins, TaHSL1 and TaHSL2, and the barley HSL protein HvHSL6A mediate TE metabolism in vitro. In the case of OsHSL4 and HvHSL6A, the TE conversion products included a derivative (metabolite 3) not seen with other HSL proteins, suggesting that the catalytic mechanism of these enzymes might differ from that of HIS1. In addition, we unexpectedly found that OsHSL1 does not recognize TE as a substrate, with TE actually acting as a weak inhibitor of OsHSL1 activity. These results revealed that it is not possible to predict the catalytic functions of HSL enzymes on the basis of only sequence similarity.

Given that the amino acid sequences of HIS1 and OsHSL1 share the highest identity (87%) among all known HSL proteins, the small differences in structure between HIS1 and OsHSL1 must determine whether TE serves as a substrate or an inhibitor. Three-dimensional structure models of HIS1 and OsHSL1 reveal differences in the putative substrate-binding pockets of these proteins (Maeda et al., 2019). On the basis of these structural models, we designed a series of mutant OsHSL1 proteins in which specific amino acids were replaced by the corresponding residues of HIS1, and we then tested the ability of these mutants to metabolize TE in vitro. Among the three residues examined, the most notable was Phe^140^ of OsHSL1, given that both HIS1 and OsHSL2 have His at this position. We therefore expected that the OsHSL1_F140H mutant might have acquired TE-metabolizing activity. However, none of the single-mutants examined (F140H, L204F, and F298L) manifested such activity. Both double-mutants containing F140H did acquire the ability to metabolize TE, and the triple-mutant showed a level of activity similar to that of HIS1. The effect of the F298L mutation was unexpected because OsHSL2 also has Phe at the corresponding position. Our results thus suggest that minor structural changes induced by amino acid substitutions in the putative substrate pocket of OsHSL1 lead to marked changes in protein function. We are currently taking a similar approach to engineer HSL enzymes for customization of their activities toward specific bTHs.

Small changes to the amino acid sequence of CYPs have been found to affect substrate specificity. Single nucleotide polymorphisms of specific CYPs have become important indicators of drug dosage appropriate for administration to individual patients. Such polymorphisms thus result in differences in drug turnover within the body and thereby influence drug efficacy (Honda et al., 2011; Ahmad et al., 2018). Our results now indicate that small sequence differences can have a substantial effect on the catalytic function of 2OGD proteins.

In conclusion, the results of the present study have revealed that 2OGD proteins, like CYPs, may play an important role in xenobiotic metabolism in plants as a result of their catalytic promiscuity.

## MATERIALS AND METHODS

### Chemicals

Daminozide [4-(2,2-dimethylhydrazinyl)-4-oxobutanoic acid] was obtained from Tokyo Chemical Industry, and both prohexadione (3,5-dioxo-4-propionylcyclohexanecarboxylic acid) and TE [ethyl-(3-oxido-4-cyclopropionyl-5-oxo) oxo-3-cyclohexenecarboxylate] were from Fujifilm Wako Pure Chemical. The plant growth regulator Primo Maxx, which contains 11.2% TE as an active ingredient, was obtained from Syngenta Japan. 3-(2-Chloro-4-mesylbenzoyl)-bicyclo[3.2.1]octane-2,4-dione (BBC-OH) was obtained from SDS Biotech. 2-{2-Chloro-4-mesyl-3-[(RS)-tetrahydro-2-furylmethoxymethyl]benzoyl}cyclohexane-1,3-dione (TFT) was also obtained from Fujifilm Wako Pure Chemical.

### Cell-Free Protein Synthesis

HIS1 and OsHSL1 proteins were prepared as described previously (Maeda et al., 2019). For preparation of other HSL proteins, the nucleotide sequences of cDNAs including the protein-coding regions were optimized for cell-free protein expression in a wheatgerm system and were then obtained as synthetic DNA molecules from Eurofins Genomics. These cDNAs included those for OsHSL2 (GenBank accession no. LC505696), OsHSL4 (LC589269), OsHSL5 (LC589270), OsHSL6 (LC589271), TaHSL1 (LC505700), TaHSL2 (LC505701), HvHSL1A (LC505697), HvHSL6A (LC505698), HvHSL6D (LC505699), SbHSL1 (LC505704), ZmHSL1A (LC505702), and ZmHSL1B (LC505703). Each cDNA was digested with the restriction enzymes SpeI and SalI, and the excised DNA fragment, including the open reading frame, was cloned into the corresponding sites of the pYT08 expression vector (Nozawa et al., 2011). Messenger RNAs for synthesis of each enzyme protein were prepared by in vitro transcription from the pYT08-based vectors, and the proteins were synthesized with the use of a wheat-germ cell-free system as described previously (Maeda et al., 2019).

### Assay of Enzyme Activity

Each substrate was dissolved in dimethyl sulfoxide to prepare a stock solution. The activity of HIS1, OsHSL1, OsHSL2, OsHSL4, OsHSL5, OsHSL6, TaHSL1, TaHSL2, HvHSL1A, HvHSL6A, HvHSL6D, SbHSL1, ZmHSL1A, or ZmHSL1B was determined by monitoring the decrease in the amount of substrate by reversed-phase HPLC with a LaChrome Elite system (Hitachi). All solutions were incubated at 30°C for 5 min prior to mixing for the enzyme reaction. The translation mixture containing each enzyme was mixed with the substrate (final concentration of 0.3 mM), FeCl2 (0.1 mM), 2OG (0.6 mM), and HEPES-KOH (100 mM, pH 7.0). The reaction mixture (final volume of 200 μl) was then incubated at 30°C for 60 min, after which the reaction was stopped by the addition of an equal volume of methanol (Nacalai Tesque) followed by incubation on ice for 15 min. The mixture was centrifuged at 20,000’ *g* for 15 min at 4°C, and the resultant supernatant was transferred to a new 1.5-ml plastic tube and dried under vacuum. The residue was dissolved in HPLC buffer, passed through a Cosmonice Filter (W) (pore size of 0.45 μm, diameter of 4 mm; Nacalai Tesque), and injected into a Pro C18 column (length of 150 mm with an i.d. of 4.6 mm, YMC), which was then subjected to elution at a flow rate of 1 ml/min at 40°C with an equal mixture of 1% acetic acid and acetonitrile. Elution was monitored by measurement of *A*284. As a negative control, the enzyme reaction was performed with a translation mixture for GFP, which did not show catalytic activity toward the tested substrates. The substrate peak for the negative control reaction was considered as the total amount of substrate, and the decrease in the amount of substrate was estimated from the peak area measured for each recombinant enzyme assay. Peak area was calculated with the use of EZChrom Elite ver. 3.1.5aJ software (Scientific Software).

### Isolation of TE Metabolites

The 200-ml reaction mixture for preparation of TE metabolites contained 20 ml of the translation mixture for HIS1 protein, TE (final concentration of 0.8 mM), FeCl2 (0.1 mM), ascorbate (1.2 mM), 20G (0.9 mM), and HEPES-KOH (100 mM, pH 7.0). The reaction mixture was incubated at 30°C for 12 h, after which the reaction was stopped by the addition of an equal volume of methanol (Nacalai Tesque) followed by incubation at 4°C for 16 h. The mixture was then centrifuged at 20,000’ *g* for 30 min at 4°C, the resulting supernatant was mixed with deionized water in order to reduce the methanol concentration to 0.3%, and the resultant solution was subjected to column purification.

A reversed-phase ODS column, Mega BE-C18 (5 g, 20 ml; Agilent), was conditioned with 100 ml of 100% methanol and equilibrated with 50 ml of deionized water. For removal of impurities, the prepared sample was applied to the column in 50ml batches, after which the column was washed with 20 ml of 10% methanol and then subjected to elution with 30 ml of 50% methanol. After confirmation that the recovered eluate contained the TE metabolites by HPLC analysis, it was transferred to a 500-ml eggplant flask, and methanol was removed with the use of a rotary vacuum evaporator in a 45°C water bath. After the addition of 100 ml of a 1:1 (vol/vol) mixture of acetonitrile and H2O, the sample was freeze-dried, and the resultant residue was dissolved in 50 ml of acetonitrile-H2O (1:1, vol/vol), transferred to a 100-ml roundbottom flask, and subjected to rotary vacuum evaporation in order to remove the acetonitrile. The resultant concentrated aqueous solution was mixed with 10 ml of 50 mM phosphoric acid, and all of the mixture was applied to a DSC-C18 (5 g/20 ml) column that had been conditioned consecutively with 20 ml of acetonitrile and 20 ml of 50 mM phosphoric acid. The flow-through fraction was discarded, the column was washed with 20 ml of a mixture of acetonitrile and 50 mM phosphoric acid (3:7, vol/vol), and the TE metabolites were eluted with 20 ml of a 4:6 (vol/vol) mixture of acetonitrile and 50 mM phosphoric acid. The eluate was recovered in a 100-ml roundbottom flask and subsequently transferred to a 200-ml separating funnel containing 100 ml of ethyl acetate, to which was then added 5 g of NaCl and 10 ml of 50 mM phosphoric acid. The mixture was subjected to extraction by shaking for 5 min at room temperature, the resulting aqueous layer was discarded, and the organic layer was recovered with 30 ml of ethyl acetate and applied to a sodium sulfate column for dehydration. The dehydrated solution was recovered in a 300-ml round-bottom flask, and ethyl acetate was removed with a rotary vacuum evaporator in a water bath (<30°C) to obtain the final dried sample.

### LC-ESI-MS

ESI-MS spectra were obtained with a Xevo G2-XS QTof MS system (Waters). The LC analysis shown in Figure 2A was performed with a Capcell Pak ADME column (150 mm, with an i.d. of 4.6 mm; Shiseido) and a mixture of acetonitrile and 0.1% acetic acid (4:6, vol/vol) at 40°C. Elution was monitored at a wavelength of 286 nm. For ESI-MS, the mode of ionization was API-ES (positive) and the signal mode was MS^E^ (50–1000 *m/z*). Fragmentor voltage was tested with a gradient range of 20 to 40 V in order to detect molecular ions and fragment ions.

### NMR Analysis

^1^H-NMR spectra (in CDCl3) and ^13^C-NMR spectra (in CDCl3) were measured with a Bruker Ascend 400 instrument at 400 and 101 MHz, respectively.

### Phylogenetic Analysis

Phylogenetic analysis for HIS1 and 13 HIS1-related sequences was performed with the use of the function “build” of ETE3 v3.0.0b32 (Huerta-Cepas et al., 2016), as implemented on GenomeNet (https://www.genome.jp/tools/ete). Alignment was performed with MAFFT v6.861b with the default options (Katoh and Standley, 2013). The tree was constructed with FastTree v2.1.8 according to default parameters (Price et al., 2009).

### Plant Growth Assay

TE-sensitivity was tested in test tubes (diameter, 2.5 cm; height, 15 cm) containing 10 ml of MS solid medium including TE (Fujifilm Wako Pure Chemical). *O. sativa* cv. Nipponbare, seeds were dehusked and surface-sterilized by two treatments with 4% sodium hypochlorite for 20 min followed by three to four rinses with sterilized water. Next, the seeds were immersed in sterilized water for two days at 30°C and germinated seeds were embedded on solid MS medium composed of 0.5 × MS salts, B5 vitamins, 3% sucrose, 0.4% gellan gum (Fujifilm Wako Pure Chemical) and TE were cultured at 27°C for 7 days with 16 hours of light (40 μmol m^-2^ s^-1^) daily. Growth inhibition rate was calculated based on the average length of the second leaf sheath.

Transgenic rice lines of *O. sativa* cv. Yamadawara, which are over-expressing *HIS1* under the cauliflower mosaic virus 35S promoter, were described in a previous report (Maeda et al., 2019). The seeds of homozygous T3 transgenic or nontransgenic rice were dehusked, surface-sterilized and embedded on the solid MS medium which contains TE (Primo Maxx), half concentration MS salts, B5 vitamins, 3% sucrose and 0.4% gellan gum. TE-sensitivity was tested in test tubes (diameter, 2.5 cm; height, 15 cm) and were cultured at 27°C for 7 days with 16 hours of light (130 μmol m^-2^ s^-1^) daily.

### Plasmid Construction for Expression of Mutant OsHSL1 Proteins

Mutations that result in amino acid substitutions in OsHSL1 were introduced into the cDNA sequence by PCR-based site-directed mutagenesis as previously described (Kanno et al., 2005). The plasmid AK241948/pFLC1, which encodes OsHSL1, was obtained from the Genebank Project of National Agriculture Research Organization (https://www.naro.affrc.go.jp/english/laboratory/ngrc/genebank_project/index.html) for use as a template. Oligonucleotide primers for the site-directed mutagenesis are listed in Supplemental Table S4. DNA fragments containing each mutated open reading frame were amplified from the pFLC1-derived plasmids by PCR with the primers HSL1-SpeF and HSL-SalR (Supplemental Table S4). The amplified fragments were digested with SpeI and SalI before cloning into the corresponding sites of pYT08, resulting in the generation of pYT08_OsHSL1_F140H, pYT08_OsHSL1_L204F, and pYT08_OsHSL1_F298L. Plasmids for the expression of OsHSL1 derivatives containing two or three amino acid substitutions were constructed as follows: pYT08_OsHSL1_F140H/L204F and pYT08_OsHSL1_F140H/F298L were prepared by site-directed mutagenesis from pYT08_OsHSL1_F140H; pYT08_OsHSL1_L204F/F298L was prepared from pYT08_OsHSL_L204F; and pYT08_OsHSL1_F140H/L204F/F298L was prepared from pYT08_OsHSL1_F140H/L204F. Each constructed plasmid was used as a template for in vitro transcription, and the resultant mRNAs were used for cell-free protein synthesis.

### Accession Numbers

Sequence data for this article can be found in the GenBank/EMBL data libraries under accession numbers: LC505696 (OsHSL2), LC589269 (OsHSL4), LC589270 (OsHSL5), LC589271 (OsHSL6), LC505700 (TaHSL1), LC505701 (TaHSL2), LC505697 (HvHSL1A), LC505698 (HvHSL6A), LC505699 (HvHSL6D), LC505704 (SbHSL1), LC505702 (ZmHSL1A), and LC505703 (ZmHSL1B).

## Supplemental Data

The following supplemental materials are available.

**Supplemental Figure S1.** Prohexadione inhibits HIS1 enzyme activity in vitro.

**Supplemental Figure S2.** MS analysis and predicted structure of a TE metabolite generated by HIS1 catalysis.

**Supplemental Figure S3.** TE weakly inhibits OsHSL1 enzyme activity in vitro.

**Supplemental Figure S4.** In vitro TE conversion assay for various HSL proteins.

**Supplemental Figure S5.** TE conversion by OsHSL1 mutant proteins in vitro.

**Supplemental Table S1.** NMR data for metabolite 1 of TE.

**Supplemental Table S2.** Effects of TE and TE-OH on the growth of rice seedlings.

**Supplemental Table S3.** Effect of *HIS1* overexpression on TE sensitivity of seedling growth in a *his1* rice mutant.

**Supplemental Table S4.** Oligonucleotide primers used in the study.

## ACKNOWLEDGMENTS

We thank S. Oyamatsu and S. Takei of Saitama University for assistance with plasmid construction and plant analysis.

